# *Pseudomonas simiae-*induced resistance in barley is subject to pathogen-dependent gene expression regulation and not associated with major changes in the phyllosphere microbiome

**DOI:** 10.64898/2026.01.25.701566

**Authors:** Anna Sommer, Samaneh Bagheri, Sanjukta Dey, Claudia Knappe, Marion Wenig, Carina Heuschmann, Susanne Kublik, Thomas Stempfl, Michael Schloter, A. Corina Vlot

**Author notes:** **Correspondence:** A. Corina Vlot.

## Abstract

Interactions of plant growth-promoting rhizobacteria with plant roots can trigger induced resistance (IR) protecting above-ground tissues from disease, a process that is increasingly recognized for its potential to support sustainable crop protection strategies. Although the molecular basis of IR has been extensively studied in *Arabidopsis thaliana*, considerably less is known about the underlying mechanisms in cereal crops. Here, we show that *Pseudomonas simae* WCS417r triggers IR in barley (*Hordeum vulgare*), reducing the propagation of *Blumeria graminis* f. sp. *hordei*, the causative agent of barley powdery mildew. Analysis of defense-associated marker genes revealed pathogen-dependent transcriptional responses during IR: *HvPATHOGENESIS-RELATED1* (*HvPR1*) was induced by powdery mildew but not by the bacterial pathogen *Xanthomonas translucens* pathovar *cerealis*, whereas *HvPR5* responded to both pathogens, with its induction by *X. translucens* depending on WCS417r-IR. To improve our molecular understanding of IR, transcriptomic responses to methyl jasmonate, salicylic acid and abscisic acid were characterized and used to identify hormone-responsive genes. Notably, several jasmonate-responsive genes were suppressed during powdery mildew infection, and this suppression was more pronounced in plants undergoing IR, indicative of priming. Finally, metabarcoding of the phyllosphere microbiome demonstrated that IR was not associated with major shifts in bacterial community composition. Alpha and beta diversity remained largely unchanged, while a limited number of amplicon sequence variants differed in abundance between treatments. Together, these results show that *P. simiae* WCS417r induces resistance against powdery mildew in barley through priming pathogen-dependent transcriptional changes while leaving the overall phyllosphere microbiome structure largely intact.

## Introduction

Induced systemic resistance (ISR) represents a crucial component of plant defense, enabling crops to enhance their innate immunity against a broad spectrum of pathogens (Pieterse *et al*., 2014; Vlot *et al*., 2021; Fiorilli *et al*., 2024; Jung *et al*., 2024). ISR is typically elicited by beneficial, soil-dwelling microorganisms colonizing the rhizosphere or the root interior. Such microorganisms can often not only elicit a state of heightened immunity but also increase resistance towards different abiotic stressors and increase plant growth; they are thus termed plant growth promoting rhizobacteria/fungi (PGPR/F). The specific protective and growth-promoting properties vary between PGPR/F and respective hosts. Root-dwelling ISR-inducing microbes include rhizobacteria such as *Pseudomonas spp, Bacillus spp. Streptomyces spp*. as well as several species of fungi, including *Serendipita indica* and *Trichoderma spp*. (Balmer *et al*., 2013; Pieterse *et al*., 2014; Weiß *et al*., 2016; Newitt *et al*., 2019; Aloo *et al*., 2022; Jung *et al*., 2024; Riseh *et al*., 2025). Because the molecular mechanisms and associated phytohormone pathways vary depending on the specific tripartite interaction between PGPR/F, plant, and challenge pathogen, we collectively summarize the elicited systemic response as induced resistance (IR) (Kesel *et al*., 2021; Vlot *et al*., 2021).

To this date, the best studied IR system is that of *Pseudomonas simiae* WCS417r in *Arabidopsis thaliana*. *P. simiae* WCS417r was one of the first PGPR described in the context of IR (Pieterse *et al*., 1996). Initially isolated from disease-supressing soil and helping to control take-all disease elicited by the fungus *Gaeumannomyces graminis* in wheat, the strain rapidly became a model organism for studying IR (Pieterse *et al*., 2021). It can elicit IR in a plethora of plants, including *A. thaliana*, eucalyptus trees, banana plants, radish, and wheat. The molecular mechanisms that underlie WCS417r-IR remain less well understood. Iron homeostasis seems to be an angling-point with the transcription factor MYB72 promoting both iron uptake in plants under iron-limiting conditions in the soil and WCS417r-IR (Zamioudis *et al*., 2014; Zamioudis *et al*., 2015). WCS417r-IR has been associated with priming of jasmonic acid (JA)-associated defense responses, promoting resistance in *A. thaliana* against pathogenic *P. syringae* pathovar tomato (*Pst*/DC3000) (Pieterse *et al*., 1998; Pozo *et al*., 2008). When interacting with *A. thaliana* leaves, WCS417r can also prime salicylic acid (SA) defense responses against *Pst*/DC3000 (Sommer *et al*., 2024).

In contrast to model systems like *A. thaliana*, IR in monocotyledonous crops, including cereals, such as barley (*Hordeum vulgare*), remains considerably less resolved (Balmer *et al*., 2013; Vlot *et al*., 2021; Shasmita *et al*., 2022; Zhao *et al*., 2024). IR in barley can be induced by a number of PGPR/F and appears mostly to be executed as a form of priming (Molitor *et al*., 2011; Duan *et al*., 2023). Systemic acquired resistance (SAR), another form of IR triggered in systemic tissues of plants undergoing a local pathogen attack, can be induced in barley using *P. syringae* pathovar *japonica* as the IR inducer (Dey *et al*., 2014). The IR response is active against pathogenic *Xanthomonas translucens* pathovar *cerealis* (*Xtc*) in a manner that appears independent of SA accumulation or signaling, whereas the same response reduces propagation of *Blumeria graminis* f.sp. *hordei* (*Bgh*), the causative agent of barley powdery mildew, in association with known SAR signaling intermediates, including pipecolic acid and volatiles (Dey *et al*., 2014; Lenk *et al*., 2019; Brambilla *et al*., 2022; Brambilla *et al*., 2023). Notably, although SAR in *A. thaliana* is classically associated with SA (Spoel & Dong, 2012; Vlot *et al*., 2021), the involvement of SA in SAR in barley appears less clear and/or at least in part pathogen-dependent (Dey *et al*., 2014; Lenk *et al*., 2018; Lenk *et al*., 2019).

Plants can actively recruit PGPR/F belowground to enhance immunity against aboveground pathogens (Hacquard *et al*., 2017; Yu *et al*., 2019). This is achieved upon the exudation of secondary metabolites from the roots of infected plants into the rhizosphere resulting in the suppression of some and enrichment of other microbiota members in the rhizosphere microbiome (Rudrappa *et al*., 2008; Stringlis *et al*., 2018). Similarly, plant immunity influences the composition of the phyllosphere microbiome of *A. thaliana* (Chen *et al*., 2020; Pfeilmeier *et al*., 2021; Sohrabi *et al*., 2023). Reciprocally, phyllosphere microbiome members can promote plant immunity against pathogens (Vogel *et al*., 2021; Liu *et al*., 2023). We have, for example, shown that propagation of WCS417r in the phyllosphere of *A. thaliana* was associated with SA-dependent IR and the enrichment of *Flavobacterium johnsoniae* in the *A. thaliana* phyllosphere microbiome (Sommer *et al*., 2024). Exposure of naïve plants to Leaf82, a *F. johnsoniae* strain from the *At-*L-Sphere collection (Bai *et al*., 2015), promoted systemic, SA-associated defense responses against *Pst*/DC3000 (Sommer *et al*., 2024).

Here, we establish the WCS417r – barley interaction as a tractable model to study PGPR-IR in cereal crops. Combining disease phenotyping with gene expression analysis and phyllosphere microbiome profiling, we show that IR and associated transcriptional responses in barley depend on both the IR-inducing PGPR and the challenge pathogen, while the composition and diversity of the phyllosphere bacterial community remain largely unaffected. By identifying phytohormone-responsive genes in barley, the data further provide new insights into barley-specific phytohormone responses during IR.

## Materials and Methods

### Plants and growth conditions

Seeds of barley (*Hordeum vulgare* L.) cultivar Golden Promise were surface-sterilized using 1.2% (v/v) sodium hypochlorite and rinsed three times with sterile water before sowing as described (Brambilla *et al*., 2023). For IR experiments, seeds were germinated and grown in the dark on ½ Murashige and Skoog medium for 3-5 days (d) prior to treatment and transfer to soil. Subsequently, plants were grown in 0.5 L pots in standard potting soil (Einheits Erde; Classic Profisubstrat, Germany) in a climate chamber (GroBank, CLF PlantClimatics GmbH, Wertingen, Germany), with a 14/10 h light/dark cycle [light intensity: ∼120 μmol cm^−2^ s^−1^ of maximum incident photosynthetically active quantum flux density levels at plant canopy] and temperatures of 20/18 °C. Plants used for phytohormone treatments were grown in a greenhouse with additional lights (HQI-TS 400W/D (Osram)) in 12-h-light/12-h-dark cycles and ∼24°C/20°C.

### IR elicitors, pathogens and treatments

IR was induced in barley using *Pseudomonas simiae* WCS417r as described (Sommer *et al*., 2021; Sommer *et al*., 2024). In short, cells were harvested from an O/N plate-grown culture (NB medium; Carl Roth, Karlsruhe, Germany) and suspended in 10 mM MgCl_2_ to a final OD_600_ of 0.2 (equalling ∼ 2 * 10^8^ colony forming units (CFU)/mL). The roots of 3-5-day-old sterile-grown barley seedlings were immersed in the bacterial suspension (or in 10 mM MgCl_2_ as control treatment) in wells of a 24-well plate for 1-2h prior to transfer of the seedlings to soil. The plants were propagated on soil for three weeks. Afterwards, leaf tissue was either harvested for molecular analysis or inoculated with *Bgh* (Swiss field isolate CH4.8) or *Xanthomonas translucens* pathovar *cerealis* (*Xtc*) (LMG7393), both of which were maintained, prepared, and inoculated as described (Dey *et al*., 2014; Lenk *et al*., 2018; Lenk *et al*., 2019; Brambilla *et al*., 2022). *Bgh* propagation was analysed at 7 days post-inoculation (dpi) using DAF-FM-DA staining to visualize the fungal hyphae and to measure relative fluorescence units quantifying the infection as described (Lenk *et al*., 2018; Lenk *et al*., 2019; Brambilla *et al*., 2022). *Xtc* propagation was analysed as described (Dey *et al*., 2014). At 4 dpi, cells were extracted from 3 leaf discs per sample in 10 mM MgCl_2_ supplemented with 0.01% (v:v) Silwet L-77 (PlantMedia, Worthington, Dublin, OH, USA); five samples were taken per biologically independent replicate experiment. For quantification, cells from each sample were serially diluted and grown on NB-plates at 28°C for two days. Subsequently, colony forming units (CFU) were counted.

Phytohormone treatments were performed in 3-4-week-old soil-grown plants in the greenhouse as described (Dey *et al*., 2014). 100 µM solutions of SA (Sigma-Aldrich), MeJA (95%; Sigma-Aldrich), and ±ABA (98%; ACROS Organics) in ddH_2_O were prepared as described (Dey *et al*., 2014) and applied to the second true leaf of the plants by syringe-infiltration. The treated leaves were harvested 24 hours post-inoculation (hpi) for RNA extraction and analysis.

### Plant phenotyping

Plant height, digital biomass, and normalized difference vegetation index (NDVI) were measured three weeks after WCS417r and control treatments using 3 dimensional (3D) multispectral imaging (Traitfinder; Phenospex B.V., Heerlen, The Netherlands). The data were analysed using the software package HortControl according to the manufacturer’s instructions (Phenospex B.V., Heerlen, The Netherlands). NDVI compares the reflectance of red and near-infrared (NIR) wavelengths as a proxy for photosynthesis with high chlorophyll content resulting in low red-light reflectance and a high reflectance of NIR. NDVI values greater than 0.6 indicate healthy vegetation with moderate to high photosynthesis.

### RNA isolation and RT-qPCR analysis

RNA for gene expression analysis was isolated from pooled samples of the second true leaf of 3-4 treated barley plants per sample using either the RNeasy Plant Mini Kit (Qiagen) or TRI Reagent (Sigma Aldrich) according to the manufacturer’s instructions. cDNA was generated using Oligo(dT) (20-mer) and SuperScript II Reverse Transcriptase (RT) (Invitrogen, Hilden, Germany) and qPCR was performed using the SensiMix SYBR Low-ROX Kit (Meridian Life Biosystems, Foster City, Ca, USA) as described (Brambilla *et al*., 2023). Transcript accumulation of *HvPR1* was analysed with primers from (Molitor *et al*., 2011) and that of the reference genes *HvEF1α* and *HvUBIQUITIN* (*HvUBI*) with primers from (Brambilla *et al*., 2022; Brambilla *et al*., 2023). Further primers used for qPCR analysis are listed in Supplementary Table S1. qPCR reactions were run in part on a 7500 real-time PCR system (Applied Biosystems) and the results were analyzed using the Fast System Software 1.3.1. qPCR reactions for Figures 2, S4, and S6 were run on a qTOWER³ Real-Time PCR system (Analytik Jena GmbH, Jena, Germany) followed by data analysis using qPCRsoft (version 4.1). Statistical analysis was done using GraphPad Prism 10 (GraphPad software, Boston, Massachusetts, USA).

### RNA-Sequencing and data analysis

Paired-end libraries of RNA isolated from phytohormone-treated plants were generated and sequenced as described (Dey *et al*., 2014). 190 Mio raw reads were generated and mapped to barley reference genome Morex v2. Adaptors were trimmed using Trimmomatic 0.39 with default settings to preserve a minimum Phred score of Q20 over 60 base-pairs. Transcript abundance was calculated using kallisto 0.46.2, based on the barley cv. Morex v2 transcript reference, available at the time of data generation. The data were analysed with R as described (Brambilla *et al*., 2022). In short, *DESeq2* was utilized to identify DEGs at a p-value threshold of <0.05 after correcting for multiple testing using the Benjamini-Hochberg-procedure (Love *et al*., 2014). Gene ontology analysis was conducted using the package *topGO* and the “weight01” algorithm using the Kolmogorov–Smirnov test. Visualisation of the data was performed using ggplot2 including the packages, *ggvenn*, *ggpubr* and *EnhancedVolcano* ((Brambilla *et al*., 2022) and reference therein). Sequences have been deposited in the SRA and are available under the accession number PRJNA1458505.

### Analysis of the phyllosphere microbiome

Microbiome analysis was performed using a metabarcoding approach with the 16S rRNA gene as marker (Sommer *et al*., 2024) in pooled samples including the second true leaf of 3-4 treated barley plants per sample. DNA was extracted using the FastPrep Soil Kit (MPbio) following the manufacturer’s instructions, with an additional leaf-grinding step using a TissueLyser (Retsch) and 1 mm glass beads (25 Hz, 2 min). The V5–V7 regions of the bacterial 16S rRNA gene were amplified from 10 ng of DNA using primers 799F and 1193R (Chelius & Triplett, 2001; Bulgarelli *et al*., 2012) and NEBNext High Fidelity Master Mix. Three PCR replicates per sample were performed (25 cycles). Amplicons were separated on 1.5% agarose gels to remove chloroplast-derived fragments, and bacterial bands were excised and purified. Replicates were quantified, pooled equimolarly, and fragment size and concentration were verified using a Fragment Analyzer. Indexed libraries were generated using Nextera XT indices, purified with magnetic beads, pooled to 4 nM, and sequenced on an Illumina MiSeq (v3, 600-cycle). Demultiplexing was performed using MiSeq Reporter Software, resulting in 2.8 mio raw reads. Sequences have been deposited in the SRA and are available under the accession number PRJNA1400912.

Amplicon data were processed using *dada2*, including quality filtering, read merging, chimera removal, and taxonomic assignment with the Silva Seeds v138 database (Yilmaz *et al*., 2014; Callahan *et al*., 2016). Reads were truncated at the first quality score ≤2 and filtered for minimum lengths of 270 bp (forward) and 150 bp (reverse), with no ambiguous bases allowed. Cyanobacteria-derived (chloroplast) ASVs and low-abundance ASVs (<0.005% of total reads) were removed. After data pre-processing, quality control and removing of chimera, the remaining sequences were assigned to 4,323 unique ASVs with 289 to 1,339 ASVs per sample. A rarefaction curve of observed ASVs showed clear saturation and thus a sufficient sequencing depth. The “empty” controls showed signs of contamination including ASVs from the bacterial standard and additionally an overrepresentation of ASV5, indicative of a possible external contamination. Consequently, the ASVs of the “empty controls” as well as ASV5 were removed from all samples prior to further analysis. Downstream analyses were performed in R using *vegan*, *phyloseq*, and *DESeq2* (Dixon, 2003; McMurdie & Holmes, 2013; Love *et al*., 2014; Schliep *et al*., 2017). Alpha diversity was calculated using Simpson’s index (SIMPSON, 1949) and β-diversity was calculated using nonmetric multidimensional scaling in R. Differentially abundant ASVs were identified using *DESeq2* (Love *et al*., 2014; Pfeilmeier *et al*., 2021).

## Results

### *P. simiae* WCS417r triggers IR in barley against *Blumeria graminis* f.sp. *hordei*

To test whether WCS417r triggers IR in barley, we dipped the roots of sterile-grown seedlings in a WCS417r suspension or in 10 mM MgCl_2_ as the mock control and transferred the plants to soil. Three weeks later, plant height, digital biomass, as well as NDVI as a proxy for photosynthesis, did not differ between WCS417r- and control-treated plants (Fig. 1A-C). This confirmed visual similarities in plant morphology between the two treatments reported in a parallel study (Rigerte *et al*., 2026). Infection of the plants with *Bgh* resulted in reduced formation of powdery mildew pustules on the leaves of WCS417r- as compared to control-treated plants (Fig. 1D). Fluorescent staining of the fungal hyphae revealed reduced fluorescence on WCS417r-treated plants as compared to the controls (Fig. 1E), and quantification of the data confirmed that WCS417r-IR significantly reduced the propagation of *Bgh* (Fig. 1F). Thus, WCS417r effectively triggered IR in barley against *Bgh* without affecting plant growth and fitness parameters prior to the infection.

**Figure 1.**
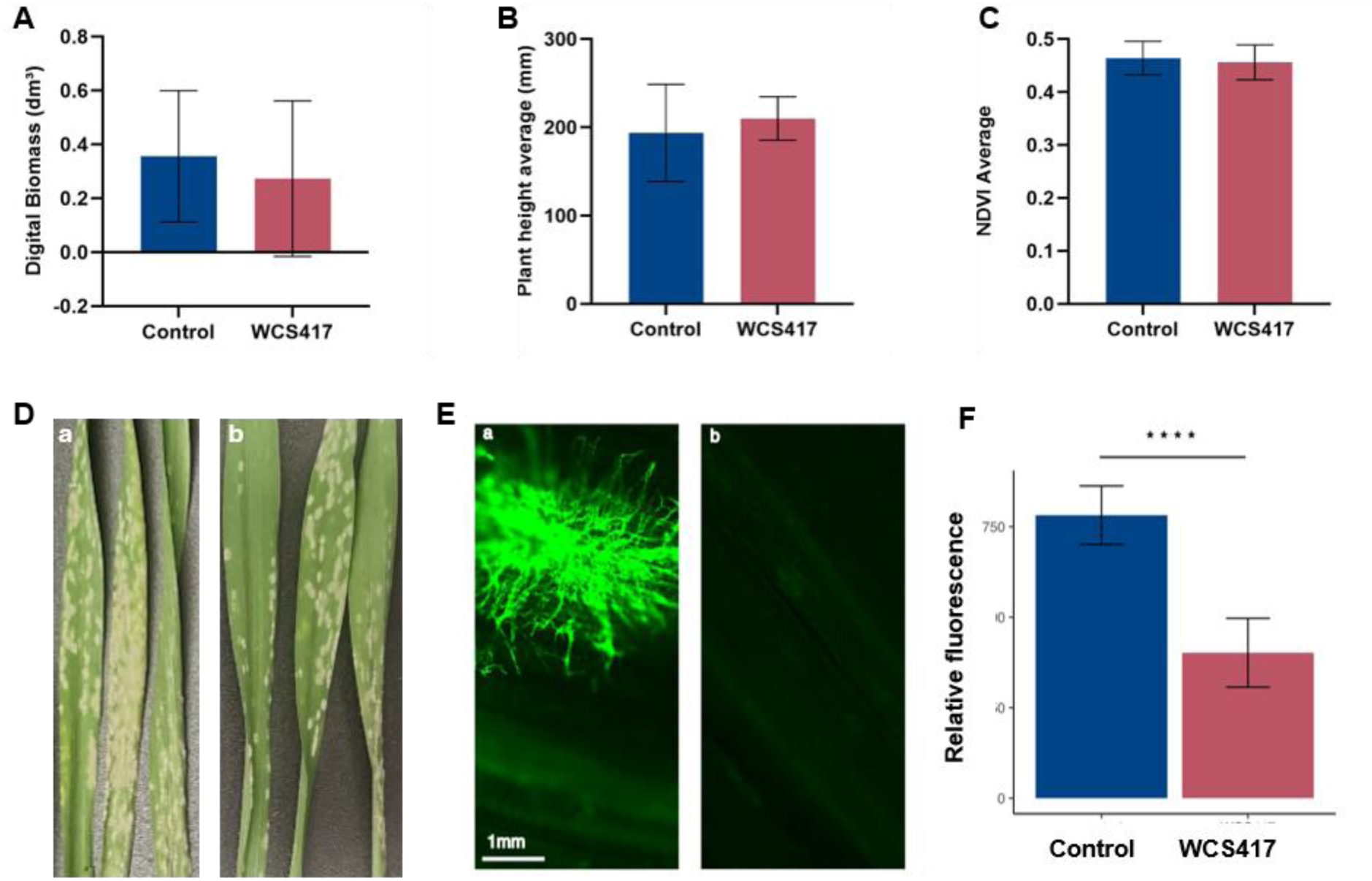
*Pseudomonas simiae* WCS417r does not significantly impact growth parameters and triggers induced resistance (IR) in barley against *Blumeria graminis* f. sp. *hordei* (*Bgh*). Sterile-grown barley seedlings (cultivar Golden Promise) were treated at the roots with *P. simiae* WCS417r (WCS417) or an appropriate control solution and transferred to soil. Three weeks later, the plants’ digital biomass (**A**), height (**B**), and Normalized Difference Vegetation Index (NDVI; **C**) were determined, and the plants were inoculated with *Bgh*. At 7 days post-inoculation (dpi), the leaves were photographed (**D**) and fungal hyphae were stained with the fluorescent dye DAF-FM-DA (**E**) and quantified by fluorescence microscopy (**F**). (**D/E**) Representative photograph of powdery mildew pustules (**D**) and microscopy images of stained *Bgh* hyphae (**E**) on control (a) and WCS417r-treated (b) plants. (**F**) Fluorescence units of *Bgh-*infected plants of the treatment groups indicated below the panel are shown relative to those of uninfected control plants. (**A-C/F**) Bars represent the mean of three biologically independent replicate experiments, each including data from 4 plants (**A-C**) or 12 leaf discs (**F**) per treatment ± SE. Asterisks indicate a statistically significant difference between the treatments (t-test ****, p <0.0001).

### Modulation of *PR* gene expression during WCS417r-IR is dependent on the challenge pathogen

In line with previous studies associating *Bgh* responses in barley with elevated *PR1* gene expression (Kogel *et al*., 1994; Molitor *et al*., 2011), we detected a strong induction of *HvPR1* transcript accumulation in response to *Bgh* in both control- and WCS417r-treated plants (Fig. 2A). Previous observations suggest that *HvPR5*, also known as *PR1A/1B* or *PWIR* (HORVU.MOREX.r3.7HG0752010), may additionally contribute to the resistance of barley against pathogens (Basak *et al*., 2024). Similarly to *HvPR1*, *HvPR5* transcript accumulation was induced by *Bgh* in both control- and WCS417r-treated plants (Fig. 2B).

**Figure 2.**
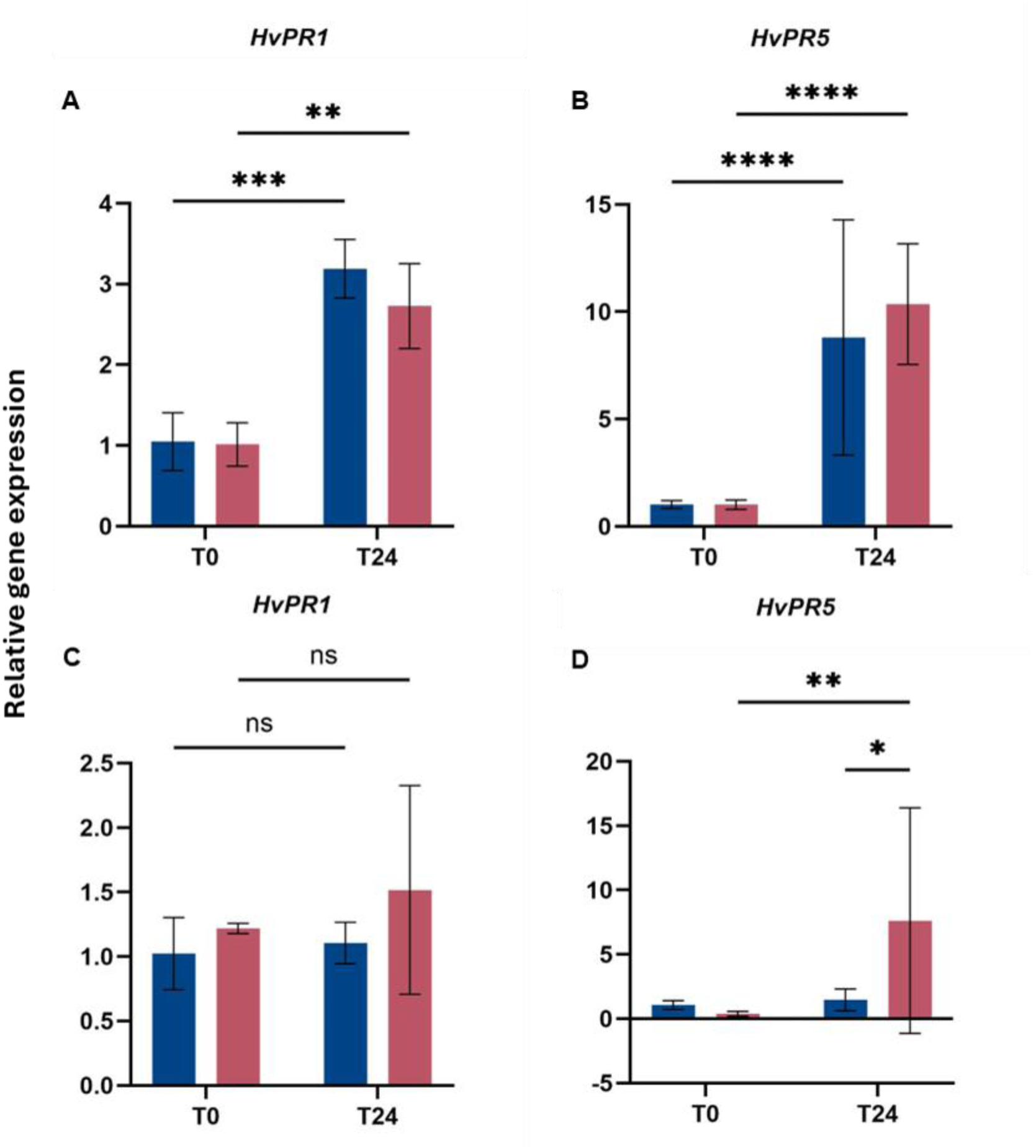
Transcript accumulation of *PR* genes in WCS417r-treated as compared to control-treated barley plants before (T0) and 24 h (T24) after infection with *Bgh* (**A/B**) or *Xanthomonas translucens* pathovar *cerealis* (*Xtc*; **C/D**). Transcript accumulation was determined by RT-qPCR of *HvPR1* (**A/C**) and *HvPR5* (**B/D**) as indicated above the panels. Data from WCS417r-treated plants are indicated in red and data from control-treated plants in blue. Transcript accumulation was normalized to that of *HvEF1α* and is indicated relative to the control at T0. Bars represent mean values +/− SD from a representative experiment, including three replicates, which was repeated three times with comparable results (**A/C**) or from three biologically independent replicate experiments, including three replicates each (**B/D**). Asterisks indicate the level of significance (Tukey’s HSD test *, p <0.05, **, p <0.01, ***, p <0.001, ****, p <0.0001; ns, not significantly different).

As introduced above, SAR in barley is effective against both *Bgh* and *X. translucens* pv. *cerealis* (*Xtc*) (Dey *et al*., 2014; Brambilla *et al*., 2022). Here, we evaluated whether WCS417r-IR was also effective against *Xtc*. In 7 biologically independent replicate experiments we detected a moderate reduction of *Xtc* propagation, which was significant in 2 experiments, in response to WCS417r-IR (Supplementary Fig. S1). In contrast to *Bgh*, *Xtc* did not appreciably change the transcript accumulation of *HvPR1* in any of three biologically independent replicate experiments that were included in the molecular analysis (Fig. 2C). However, *HvPR5* transcript accumulation was induced by the *Xtc* challenge infection in WCS417r-treated plants but not in the controls (Fig. 2D). Thus, although WCS417r-IR against *Xtc* was not robust, the WCS417r treatment primed the induction of *HvPR5* in response to *Xtc.* Moreover, the overall transcriptional response of *HvPR1* and *HvPR5* differed depending on the challenge pathogen.

### Barley transcriptional responses to MeJA, ABA, and SA

While *HvPR1* was induced by *Bgh,* the same gene did not appear sensitive to SA (Supplementary Fig. S2) or its functional analog 2,6-dichloro-isonicotinic acid (Beßer *et al*., 2000), confirming our previous observations that phytohormone associations with IR in barley appear to differ from those in *A. thaliana* (Dey *et al*., 2014). For this reason, we set out to assess the barley transcriptional response to the phytohormones SA, JA, and ABA with the aim of identifying phytohormone-responsive genes in barley. We treated greenhouse-grown barley plants with 100 µM SA, MeJA, or ABA by leaf infiltration and analysed the transcriptional response in comparison to a water-treated control group by RNA-sequencing of samples taken 24 hpi.

In MeJA-treated plants, we found 695 DEGs (400 upregulated, 295 downregulated). RNAseq analysis of ABA-treated plants revealed 116 DEGs (44 upregulated, 72 downregulated), while SA treatment resulted in 149 DEGs (77 upregulated, 72 downregulated) (Supplementary Table S2). Among these, 3.31% were shared between the responses to MeJA and SA, 5.14% between the responses to MeJA and ABA, and 0.46% between the responses to SA and ABA, while 0.34% were common between all treatments (Supplementary Fig. S3). Barley transcriptional responses to MeJA, ABA, and SA thus appeared mostly distinctive between phytohormones. To gain further insight into the characteristics of the DEGs with the highest log_2_-fold change per treatment, we utilized the tool Monocot PLAZA5.0 (van Bel *et al*., 2022) to examine predicted gene functions (Supplementary Table S2); gene names introduced below were verified and/or predicted based on the results of basic local alignment search tool (BLAST) using NCBI protein-BLAST (McGinnis & Madden, 2004) and protein sequences derived from Monocot PLAZA5.0.

Among the 20 DEGs with the highest log_2_-fold change under MeJA treatment, three were downregulated and 17 upregulated (Supplementary Table S2). The latter included a *JASMONATE-INDUCED PROTEIN* (Horvu_MOREX_3H01G721200), which was also responsive to SA, as well as a possible *JASMONATE-INDUCED OXYGENASE2/4-like* (*HvJOX2/4-like*; Horvu_MOREX_4H01G310300) and an acetyl-coenzyme A synthetase predicted to be involved in glucosinolate biosynthesis. Notably, the upregulated genes further included four genes that were predicted to be responsive to ABA. The downregulated genes included an *EXPANSIN* which was also downregulated among the 20 DEGs with the highest log_2_-fold change under ABA treatment, suggesting overlaps between the barley response to MeJA and predicted ABA responses in plants.

Among the 20 DEGs with the highest log_2_-fold change under ABA treatment were 12 downregulated and 8 upregulated genes. Among the downregulated genes were six which were associated with responses against abiotic and/or biotic stress, including three peroxidases and a receptor-like kinase predicted to act as a pattern recognition receptor of the bacterial elicitor flagellin22. The upregulated genes included one *DEHYDRIN* and a *PROTEIN PHOSPHATASE 2C*, both classically associated with ABA signaling (Raghavendra *et al*., 2010).

Among 20 DEGs with the highest log_2_-fold change under SA treatment, 19 were upregulated and one was downregulated (Supplementary Table S2). In addition to the *JASMONATE-INDUCED PROTEIN* shared with the response to MeJA, SA induced at least one further predicted JA-responsive gene, *ACID PHOSHATASE 1*, with predicted roles in plant responses to wounding and insect attack. Six further upregulated genes were associated with defense responses to biotic stress, including three DEGs with predicted roles in SA metabolism or signaling. Notably, additional five upregulated genes were associated with the process of translation.

Overall, there is a recognizable trend among the 20 top regulated genes in response to the three phytohormones tested with responses to abiotic stress dominating the responses of barley to MeJA and ABA, while biotic stress tolerance more prominently appeared associated with the barley response to SA. At the same time, classical marker genes, such as *PR1* for SA (van Loon *et al*., 2006), may not prove useful to link barley transcriptional responses to a particular pathway. For this reason, we set out to identify genes that are responsive to MeJA, ABA, and SA specifically in barley. Genes were selected from those DEGs, which were upregulated at least ∼4-fold (or 2 log_2_), and had a mean read count of at least 40 (Supplementary Table S2). To confirm a reliable upregulation of the target genes after phytohormone-treatment, RT-qPCR analysis was performed on samples from additional, biologically independent phytohormone treatment assays.

To detect MeJA-dependent gene induction, we monitored the transcript accumulation of the genes Horvu_MOREX_7H01G720100 encoding a predicted GDSL lipase (*HvGDSL-like*), Horvu_MOREX_5H01G408900 (*HvABC TRANSPORTER G FAMILY MEMBER 1-like*; *HvABCG1-like*), and Horvu_MOREX_4H01G310300 (*HvJOX2/4-like*). To detect ABA-related transcript accumulation, we tested primers for the genes Horvu_MOREX_3H01G553800 (*HvDEHYDRIN3-like*; *HvDNH3-like*), Horvu_MOREX_6H01G028000 (*HvSUCROSE:SUCROSE FRUCTOSYLTRANSFERASE*; *HvSST-like*), and Horvu_MOREX_2H01G364600 (Photosystem II complex component PsbR; *HvPSBR*). For SA-induced gene expression patterns, we established primers for the genes Horvu_MOREX_5H01G376100 (*HvWKRY76-like*), Horvu_MOREX_2H01G572800 (*HvDMR6-like OXYGENASE1-like*; *HvDLO1-like*), and Horvu_MOREX_3H01G001000 (transcription factor *HvbHLH167-like isoform X2*; *HvbHLH 167-like*). In three independent biological replicate experiments, the transcript accumulation of the selected genes increased in response to the respective phytohormone treatments by at least 2.5-fold and averaged 5.7-fold across all treatments and transcripts tested (Fig. 3).

**Figure 3.**
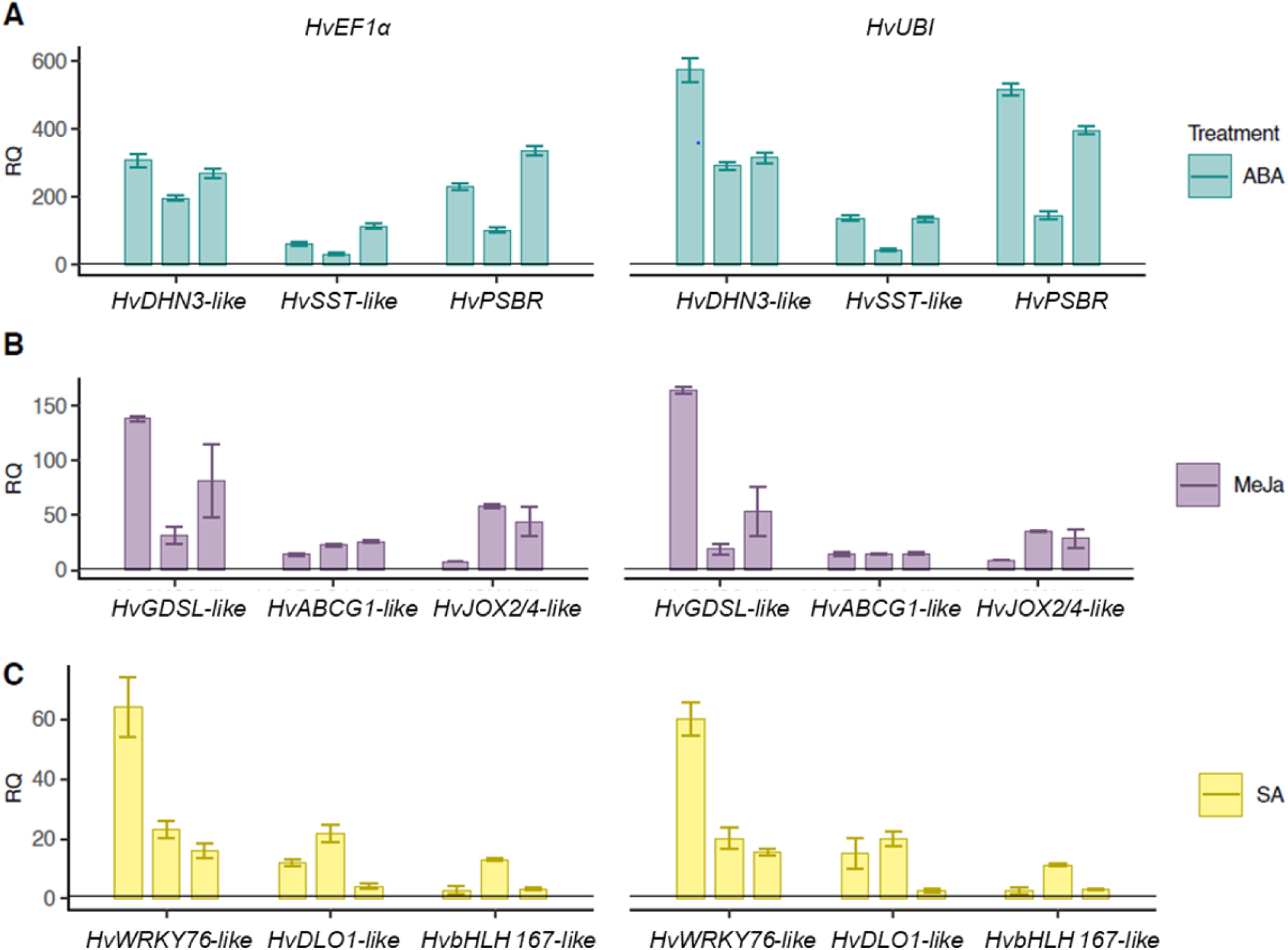
RT-qPCR validation of phytohormone-responsive DEGs. Relative accumulation is shown of ABA-responsive (**A**), MeJA-responsive (**B**), and SA-responsive (**C**) transcripts as indicated below the panels. Transcript accumulation in the panels on the left was normalized to that of *HvEF1α* and in the panels on the right to that of *HvUBIQUITIN* (*HvUBI*) as indicated at the top of the figure. Transcript accumulation is shown relative to that in appropriate water-treated controls which are indicated by black lines in the panels. Bars represent average values +/− SD from three technical replicates; the results from three biologically independent replicate experiments are shown in individual bars for each gene per treatment.

### IR against *Bgh* may be associated with reduced jasmonate signaling in barley

To gain a better understanding of the WCS417r-IR mechanism, we used the genes established above to test for phytohormone-associated gene expression changes during WCS417r-IR. Of the tested genes, all three MeJA-responsive genes displayed a trend towards reduced transcript accumulation in WCS417r- as compared to control-treated plants before infection (T_0_ in Fig. 4). Because statistical analysis using Shapiro-Wilk rejected a normal distribution of the data and fold changes varied among biologically independent replicate experiments (see Supplementary Fig. S4 for an example), a WCS417r-IR-associated reduction of MeJA-associated transcripts remained below the threshold for statistical significance. Further, the transcripts of three genes were reduced in response to *Bgh* associating their transcript accumulation with the barley response to infection. These genes included the ABA-responsive *HvDHN3-like* and the SA-responsive genes *HvDLO1-like* and *HvbHLH167-like* (Fig. 4). Notably, two of the MeJA-responsive genes, *HvGDSL-like* and *HvJOX2/4-like*, also displayed reduced transcript accumulation in response to *Bgh* while this was significant in control-treated but not in WCS417r-treated plants (Fig. 4). Transcripts of the MeJA-responsive gene *HvABCG1-like* went on to display a typical priming pattern. The already reduced transcript accumulation in response to the WCS417r treatment appeared to be followed by a further reduction in response to the following *Bgh* infection (Fig. 4). This was mirrored by similar transcript accumulation patterns of *HvJOX2/4-like*, but remained an insignificant trend for both transcripts, possibly due to the prior insignificant reduction of transcripts in WCS417r-treated plants prior to infection (Fig. 4 and Supplementary Fig. S4).

**Figure 4.**
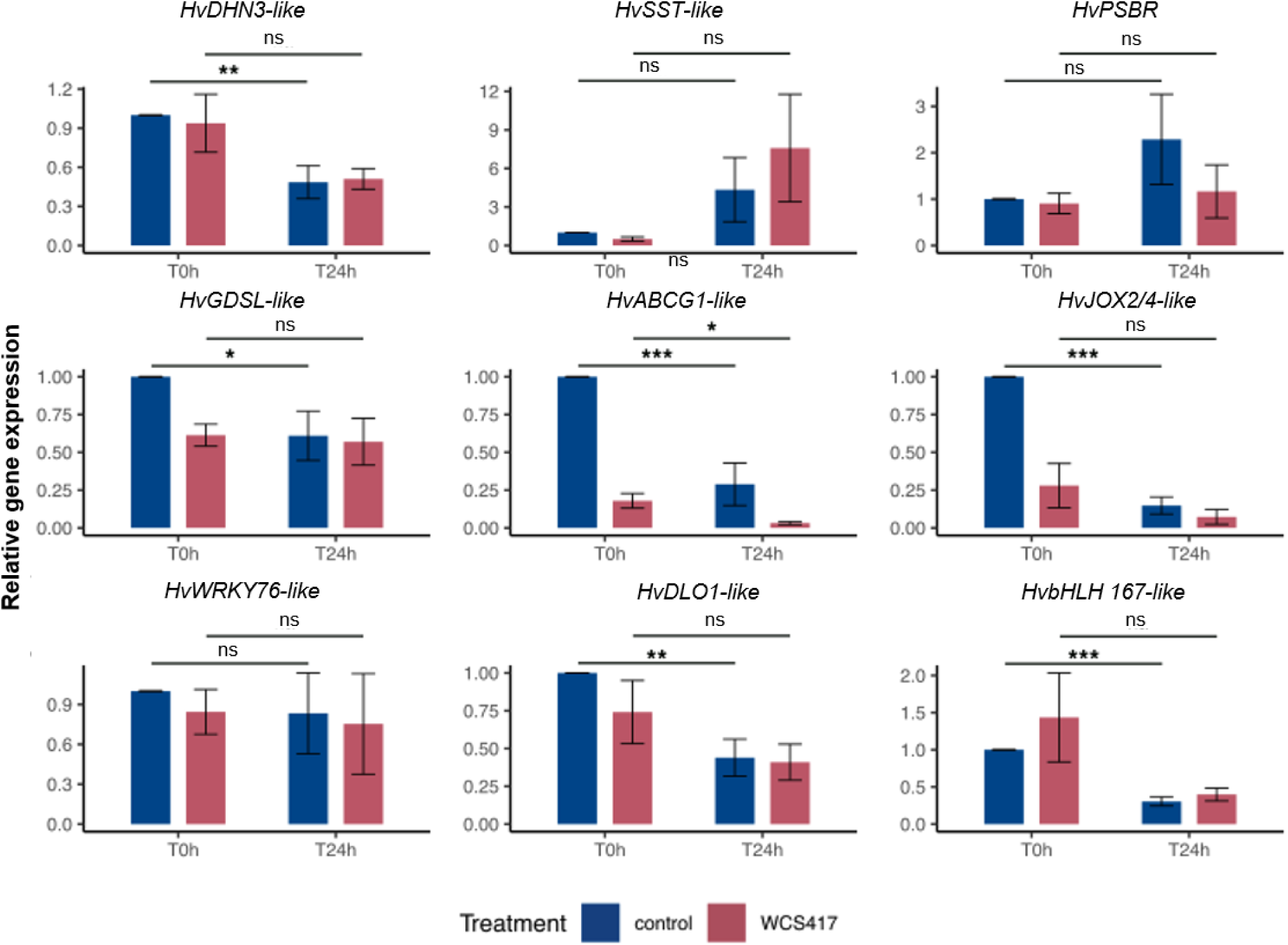
Relative transcript accumulation of phytohormone-responsive genes analysed by RT-qPCR in samples from WCS417r-treated as compared to control-treated barley plants before (T0h) and 24 h after infection of the plants with *Bgh* (T24h). Upper panels: ABA-responsive genes, middle panels: MeJA-responsive genes, and bottom panels: SA-responsive genes, as indicated above the panels. Transcript accumulation was normalized to that of *HvEF1α* and is shown relative to the control at T0. Bars represent mean values +/− SE from three biologically independent replicates, asterisks indicate level of significance (Tukey’s HSD test *, p <0.05, **, p <0.01, ***, p <0.001, ****, p <0.0001; ns, not significantly different).

Although the responsiveness of genes to phytohormones may heavily depend on the experimental conditions, phytohormone concentration, and treatment duration, it appears striking that all three MeJA-responsive genes displayed comparable transcript accumulation patterns during WCS417r-IR against *Bgh.* Therefore, we aimed to explore whether this may represent a general IR-associated response. To this end, plants were treated with *Bacillus thuringiensis* (*Bt*), previously shown to trigger IR in plants (Seo *et al*., 2012; Akram *et al*., 2013; Hyakumachi *et al*., 2013; Takahashi *et al*., 2014; Sommer *et al*., 2021). Subsequent inoculation of the plants with *Bgh* resulted in the formation of fewer powdery mildew pustules on the leaves of *Bt-*treated plants as compared to the controls (Fig. 5A). This was accompanied by reduced fluorescence upon staining of the fungal hyphae (Fig. 5B) and quantification of the data confirmed that *Bt*-IR significantly reduced the propagation of *Bgh* in barley (Fig. 5C). Both *HvABGC1-like* and *HvJOX2/4-like* displayed a comparable reaction to *Bgh* in plants undergoing IR triggered by WCS417r and *Bt* (Fig. 5D). Together, the data suggest that at least a subset of JA-sensitive genes is subject to downregulation upon exposure of barley plants to two different PGPR. This downregulation may be exacerbated by a subsequent challenge infection of the plants with *Bgh*, suggesting a priming effect associated with IR in barley against *Bgh*.

**Figure 5.**
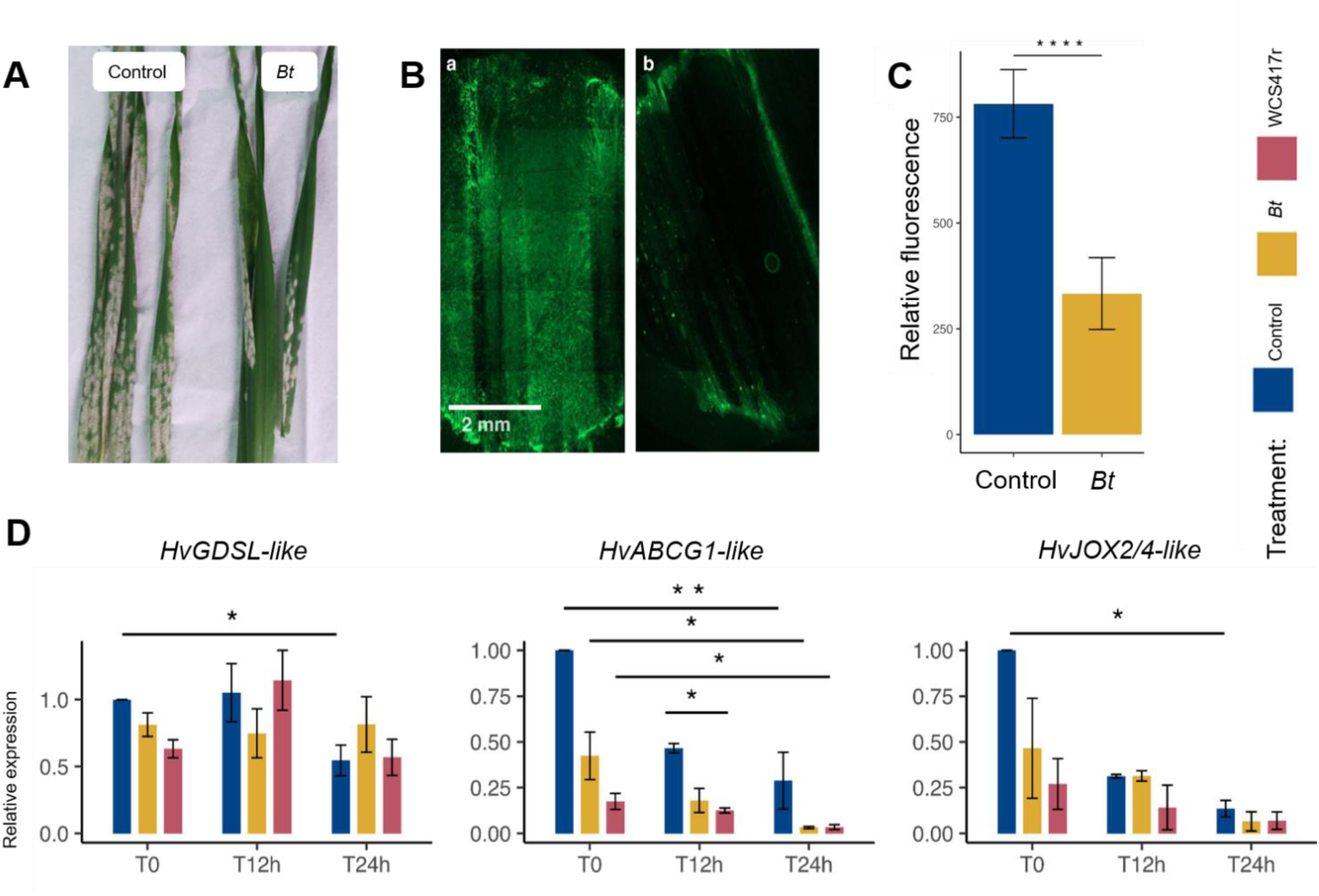
*Bacillus thuringiensis* (*Bt*) triggers IR in barley against *Bgh*. Sterile-grown barley seedlings were treated at the roots with *Bt* or an appropriate control solution and transferred to soil. Three weeks later, the plants were inoculated with *Bgh*. At 7 days post-inoculation (dpi), symptoms were photographed (**A**) and the fungal hyphae were stained with the fluorescent dye DAF-FM-DA and quantified by fluorescence microscopy. (**B**) Representative microscopy images of stained *Bgh* hyphae on control (a) and *Bt*-treated (b) plants. (**C**) Fluorescence units of *Bgh-*infected plants of the treatment groups indicated below the panel are shown relative to those of uninfected control plants. Bars represent the mean of three biologically independent replicate experiments, each including data from 12 leaf discs per treatment ±SE. Asterisks indicate a statistically significant difference between the treatments (t-test ****, p <0.0001). (**D**) Relative transcript accumulation of selected MeJA-responsive genes as indicated above the panels, analysed by RT-qPCR in samples from *Bt-* (in yellow) or WCS417r-treated (in red) as compared to control-treated barley plants (in blue) before (T0h) and 12 h (T12h) or 24 h (T24h) after infection of the plants with *Bgh*, as indicated below the panels. Transcript accumulation was normalized to that of *HvEF1α* and is shown relative to the control at T0. Bars represent mean values +/− SE from three biologically independent replicates, asterisks indicate level of significance (Tukey’s HSD test *, p <0.05, **, p <0.01).

### Phyllosphere microbiome responses to IR in barley

We previously reported that WCS417r-IR has a significant impact on the microbiome of the phyllosphere in *A. thaliana* (Sommer *et al*., 2024). Therefore, we examined if a similar connection between IR and the phyllosphere microbiome also exists in barley. To do so, we analysed 16S rRNA gene amplicon sequences in the phyllosphere of WCS417r- as compared to *Bt-* and control-treated plants (Supplementary Table S3). First, we checked for the presence of the IR-eliciting strains in the microbiome of the barley phyllosphere. To this end, we filtered the ASVs of all samples for the genus *Pseudomonas* and *Bacillus*, and blasted the obtained sequences using NCBI Blast (McGinnis & Madden, 2004). None of the Pseudomonas-derived sequences matched that of the 16S rRNA gene of *Pseudomonas simiae*, indicating that WCS417r did not migrate to the phyllosphere upon treatment of barley. However, since sequence reads matching the 16S rRNA gene of *Bt* were equally distributed across all three treatment groups, we focussed further analyses on the WCS417r treatment in comparison to the control only.

Among the ten most abundant phyla found in the barley leaf microbiome were Proteobacteria, Firmicutes as well as Actinobacteriota (Fig. 6A). In addition, we found Verrumicrobiota, Myxococca, Fibrobacteriota, Deinococcota, Bdellovibrionota as well as Acidobacteriota. Next, we determined if WCS417r-IR treatment introduced changes in the absolute number of detected ASVs present per sample as a measure of species richness as well as the associated alpha-diversity (SIMPSON, 1949). We did not find significant differences in the observed numbers of ASVs or diversity with respect to treatment (Fig. 6B). Principal component analysis as well as nonmetric multidimensional scaling of weighted Unifrac-distances (Lozupone & Knight, 2005) also did not yield clear clustering according to treatment or replicate, excluding changes in β-diversity (Fig. 6C and Supplementary Fig. S5).

**Figure 6.**
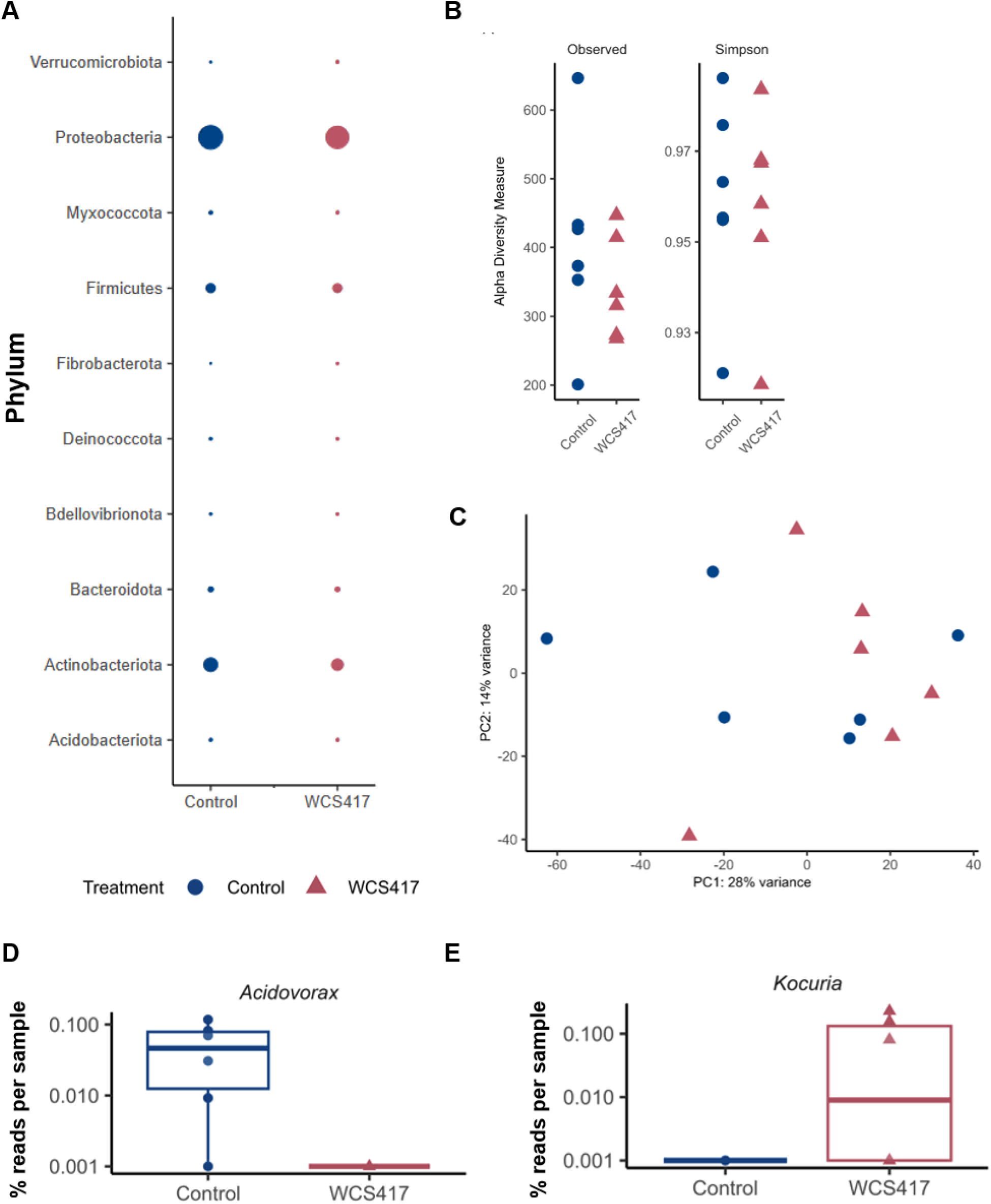
Microbiome of barley plants analysed by 16S rRNA gene amplicon sequencing in samples from WCS417r-treated (in red) as compared to control-treated barley plants (in blue); data are derived from six biologically independent replicate experiments. (**A**) 10 most abundant phyla represented by amplicon sequence variants (ASVs) in the respective treatment groups; bubble sizes correlate with the number of ASVs per phylum. (**B**) Species richness according to the observed absolute number of ASVs per treatment (left panel) and the associated alpha diversity according to the Simpson’s index (right panel). (**C**) Principal component analysis of the ASV composition per sample. (**D/E**) Relative abundance of ASV340 (**D**) and ASV316 (**E**) in the barley phyllosphere. Boxplots indicate average number of sequenced reads from six samples in percentage of reads per sample ± min and max values; individual data points are indicated.

Finally, we mined the data for ASVs which were significantly differentially abundant depending on the treatment. In the comparison group WCS417r vs. control, we found 17 ASVs with a significantly different abundance in the two data sets (Supplementary Table S4). Among the differentially accumulating ASVs was ASV340, which was present in 5 out of 6 control samples but not detected in samples from WCS417r-treated plants, thus displaying a possible depletion in WCS417r-treated plants (Fig. 6D). On the other hand, several ASVs appeared to be enriched in WCS417r-treated in comparison to control-treated plants. Among them were ASV118 of the species *Serratia marcescens* and ASV316 of the species *Kocuria rhizophila* (Fig. 6E). Thus, while the overall species richness and diversity in the phyllosphere microbiome of barley does not appear to change in response to a WCS417r-IR treatment, the relative abundance of specific microbiota may be subject to moderate shifts.

## Discussion

In this work, we established the barley – *P. simiae* WCS417r interaction as a model IR system promoting defense against powdery mildew caused by *Bgh* in barley. In a parallel study, ASVs corresponding to WCS417r were detected among 16S amplicon sequence reads of rhizosphere samples from the treated plants, suggesting that the bacteria remained present in the rhizosphere until harvest (Rigerte *et al*., 2026). Whereas it remains possible that WCS417r did not effectively colonize barley roots, exposure of the roots to the bacteria enhanced the resistance of the leaves against *Bgh* (Fig. 1). Notably, WCS417r-IR was not accompanied by detectable accumulation of WCS417r in the barley phyllosphere, suggesting that IR was established as a classical ISR response depending on plant-mediated processes priming shoot tissues for enhanced resistance against pathogens (Pieterse *et al*., 2014; Vlot *et al*., 2021).

PGPR-IR protects *A. thaliana* from both biotrophic and necrotrophic pathogens and this is associated with SA and JA responses (reviewed in (Pieterse *et al*., 2014; Schenk & Schikora, 2014; Vlot *et al*., 2021). Many studies focus on gene expression and priming responses in individual PGPR-plant-pathogen systems, sometimes differentiating between responses to different PGPR (e.g. (Duan *et al*., 2023) but not necessarily between challenge pathogens. In *A. thaliana,* WCS417r-IR enhances the resistance of the plants against hemi-biotrophic *Pseudomonas syringae* bacteria and this is associated with priming of jasmonate responses (Pozo *et al*., 2008). WCS417r-IR also protects *A. thaliana* from necrotrophic pathogens and both responses appear to converge on iron-deficiency (Romera *et al*., 2019; Trapet *et al*., 2021). Although plant defense mechanisms against biotrophic and necrotrophic pathogens differ and may even act antagonistically (Pieterse *et al*., 2012; Spoel & Dong, 2012; Broekgaarden *et al*., 2015), studies often focus on similarities between the mechanisms downstream of IR against biotrophic and necrotrophic pathogens arguing that broad-spectrum immunity may result from circumventing antagonistic crosstalk between SA and JA (van Wees *et al*., 2000; Spoel *et al*., 2007; Schenk & Schikora, 2014; Wittek *et al*., 2015; Trapet *et al*., 2021). Here, we show that WCS417r primes barley for enhanced *HvPR5* transcript accumulation when inoculated with *Xtc* (Fig. 2), while the same pathogen does not affect *HvPR1* transcript accumulation. Although priming of *HvPR5* did not appear to be associated with robust defense of barley against *Xtc*, the same gene transcript was induced along with *HvPR1* by *Bgh.* Interestingly, a possible downregulation of *HvJOX2/4-like* in response to WCS417r-IR and *Bgh* may also be driven largely by the challenge pathogen. In contrast to *Bgh,* which reduces *HvJO2/4-like* transcript accumulation, *Xtc* induces *HvJOX2/4like* and this induction is compromised in plants undergoing WCS417r-IR (Figs. 4 and 5 and Supplementary Fig. S6). Taken together, transcriptional responses downstream of IR and infection vary depending on both the IR trigger and the challenge pathogen, and this may at least in part drive broad-spectrum immunity adapting primed gene expression changes to specific challenge pathogens.

Until now, defense in barley against powdery mildew has been associated with elevated SA accumulation and/or responses (Torres *et al*., 2017; Laupheimer *et al*., 2024). While exogenous SA enhances the resistance of barley against powdery mildew (Lenk *et al*., 2018), SA does not appear to be essential for early (penetration) resistance responses against the causative *Bgh* fungus (Huckelhoven *et al*., 1999; Jain *et al*., 2004; Keisa *et al*., 2011). Additionally, jasmonate responses have been associated with resistance in barley against *Bgh* (Walters *et al*., 2002; Xu *et al*., 2022; Krasauskas *et al*., 2024). Studies have shown transient upregulation of jasmonate accumulation and/or signaling during early stages of a *Bgh* infection, returning to basal levels or below at later infection stages, as well as a moderate induction of resistance to *Bgh* in response to exogenous MeJA (Sang *et al*., 2021; Xu *et al*., 2022; Krasauskas *et al*., 2024). Together, the data suggest that JA may promote early defences against *Bgh* while its subsequent downregulation may promote enhanced resistance, for example associated with SA, against *Bgh* at later infection stages. In this study, although *JOX* gene products inactivate JA, effectively downregulating JA signaling (Caarls *et al*., 2017), the observed downregulation of JA-responsive genes, including *HvGDSL-like*, *HvABCG1-like*, *HvJOX2/4-like,* and at least one *JAZ* suggests a possible downregulation of JA responses at 24h after the inoculation of barley with *Bgh* (Fig. 4 and Supplementary Table S2).

Typically, priming is believed to exacerbate the differential regulation of infection-responsive transcripts early after a challenge inoculation, with responses in infected control plants catching up over time (Conrath *et al*., 2015; Martinez-Medina *et al*., 2016; Mauch-Mani *et al*., 2017; Zeier, 2021). For this reason, primed gene expression changes are often studied within a short time frame after a challenge inoculation (Molitor *et al*., 2011; Lee *et al*., 2012; Bernsdorff *et al*., 2016). In barley, *Serendipita indica*-induced resistance primes an early upregulation of *HvPR1* as detected 12h after a *Bgh* challenge inoculation (Molitor *et al*., 2011). In the current study, the downregulation of *HvABCG1-like* transcripts was significantly more pronounced, and thus primed, in WCS417r-treated than in control-treated plants at 12h after the *Bgh* challenge inoculation (Fig. 5D). A similar trend was observed when analysing transcript accumulation of *HvJOX2/4-like*. The JA-responsive gene *HvGDSL-like* is both upregulated in response to MeJA and downregulated by ABA (Supplementary Table S2). The latter may explain why its response to *Bgh* remained less pronounced as compared to that of *HvABCG1-like* and *HvJOX2/4-like* (Figs. 4 and 5). Because a consistent response of ABA-responsive genes was not observed, the data suggest that JA responses are at least in part downregulated at 24h after the *Bgh* inoculation and that this response is exacerbated by WCS417r-IR and thus primed. In that respect, it is of interest to note that the observed induction of *HvJOX2/4-like* transcript accumulation in response to *Xtc* appears to be compromised in plants undergoing WCS417r-IR (Supplementary Fig. S6). In future, it will be of interest to define the role of JA signaling in early defense of barley against *Bgh* to mitigate whether the compromised JA response observed here is part of the endogenous resistance response of barley against powdery mildew caused by *Bgh* or reflects a general primed response dependent on WCS417r-IR.

The plant immune status appears to take a notable effect on the composition of the phyllosphere microbiota with *A. thaliana* defense-compromised mutants displaying significant shifts in their phyllosphere microbiome (Chen *et al*., 2020; Pfeilmeier *et al*., 2021; Pfeilmeier *et al*., 2024). We previously showed that WCS417r-IR in *A. thaliana* was accompanied by the migration and propagation of WCS417r and the recruitment of a beneficial *Flavobacterium* sp. to the phyllosphere, likely due to microbe – plant – microbe interactions depending on both WCS417r and the plant immune status (Sommer *et al*., 2024). In the current study, WCS417r was not detected in the phyllosphere of barley, indicating the induction of ISR in the classical sense (Pieterse *et al*., 2014) in WCS417r-treated barley. At the same time, WCS417r-IR priming did not cause a significant shift in the phyllosphere microbiome of barley (Fig. 6) largely uncoupling primed immunity from microbiome dynamics in the phyllosphere of barley. Nevertheless, 17 ASVs displayed differential accumulation in the phyllosphere of WCS417r- compared to control-treated barley (Supplementary Table S4). Among them, we detected the depletion of the bacterial leaf blight pathogen *Acidovorax avenae* (Schaad *et al*., 2008). At the same time, two possible beneficial strains appeared enriched. *S. marcescens* has been described as a PGPR with growth-promoting properties (Zheng *et al*., 2022; Zhang *et al*., 2024), providing protection against cadmium stress and various plant diseases, including potato late blight caused by *Phytophtora infestans* and rice sheath blight caused by *Rhizoctonia solani* (Khan *et al*., 2017; El-Esawi *et al*., 2020; Jiang *et al*., 2025; Liu *et al*., 2025). *Kocuria rhizophila* is an endophytic bacterium which has been reported to display plant growth-promoting properties in different plants, for example enhancing salt and drought stress tolerance in wheat and tomato (Afridi *et al*., 2019; Mauceri *et al*., 2024; Dargiri & Samsampour, 2025). While this seems reminiscent of reported recruitment of beneficial microbes to the rhizosphere of diseased plants (Berendsen *et al*., 2018; Stringlis *et al*., 2018), it remains unclear whether the minor changes detected here take an effect on plant fitness and/or are notable in the vast abundance of microbiota that make up the phyllosphere microbiome.

In conclusion, this study shows that *P. simiae* WCS417r triggers IR in barley at least in part through priming gene expression changes that respond to both the priming trigger and the challenge pathogen. The phyllosphere microbiome did not appear to significantly react to WCS417r-IR priming, supporting the potential for application of PGPR-IR as it enhances resistance without changing the plant ecosystem.

## Author Contributions

ACV conceptualized and AS, MS, and ACV planned the project; MS and ACV acquired funding; AS, SB, SD, TS, and MS designed methodologies; AS, SB, SD, CK, MW, CH, and SK performed experiments; AS, SB, CK, MW, SK, and TS analysed the data; AS and ACV wrote the first draft of the manuscript; MS, SB, and ACV edited the manuscript and all authors reviewed and agreed to the content of the work.

## Acknowledgements

The authors are grateful to Dr. Michael Rothballer (Helmholtz Munich, Germany) for providing *Bacillus thuringiensis*, to Dr. Kathrin Paulus-Tremel (University of Bayreuth, Germany) for technical support and helpful discussion, and to Dr. Klaus F.X. Mayer and his team (Helmholtz Munich, Germany) for mapping the RNA-Seq data derived from barley treated with different phytohormones and for helpful discussions.

## Supplementary Material

**Supplementary Table S1** Oligonucleotides used for qPCR

**Supplementary Table S2** RNA-sequencing results summarizing differentially expressed genes (DEGs) in barley in response to phytohormones

**Supplementary Table S3** ASV counts in WCS417r-, *Bt-*, and control-treated plants derived from 16S rRNA gene amplicon sequencing

**Supplementary Table S4** Relative abundance of distinct ASVs analysed by 16S rRNA gene amplicon sequencing in samples from WCS417r-treated as compared to control-treated barley plants.

**Supplementary Figure S1** Relative transcript accumulation of *HvPR1* in response to phytohormones

**Supplementary Figure S2** WCS417r-IR against *Xanthomonas translucens* pathovar *cerealis* (*Xtc*)

**Supplementary Figure S3** VENN diagramm of phytohormone-responsive differentially expressed genes (DEGs) in barley.

**Supplementary Figure S4** *HvJOX2/4-like* transcript accumulation during WCS417-IR against *Bgh*

**Supplementary Figure S5** Nonmetric Multidimensional scaling (NMDS) of microbial composition per treatment and replicate.

**Supplementary Figure S6** *HvJOX2/4-like* transcript accumulation in response to *Xtc* in plants undergoing WCS417r-IR.

## Data Availability Statement

Data derived from RNA-sequencing and from 16S rRNA gene sequencing have been deposited in the SRA and are available under the accession numbers PRJNA1458505 (RNA-seq phytohormone treatments) and PRJNA1400912 (16S amplicon sequencing). All other data are available from the corresponding author upon request.

