## Supplementary Tables and Figures for "*Pseudomonas simiae-*induced resistance in barley is subject to pathogen-dependent gene expression regulation and not associated with major changes in the phyllosphere microbiome"

| Gene | Locus ID | Locus (Ensembl Plants) | Primer sequences 5’ → 3’ | F/R |
|---|---|---|---|---|
| *HvDHN3-like* | Horvu_MOREX_3H01G553800 | LOC123440604 | CGGGAGGTCACAAGAACGG | F |
| | | | AGTGGTGTTCCCATCCATTCC | R |
| *HvSST-like* | Horvu_MOREX_6H01G028000 | LOC123401241 | ACCCTTTGCTCGTCAACTGG | F |
| | | | TCGTCGGAGCCGTCAAAC | R |
| *HvPSBR* | Horvu_MOREX_2H01G364600 | LOC123426277 | ACGGAGCAAATGTTGATGGG | F |
| | | | TAGCCACCCAGAGCAACAAC | R |
| *HvWKRY76-like* | Horvu_MOREX_5H01G376100 | LOC123399557 | CCCATCACCAACGATGAGGC | F |
| | | | TGGTTGCTGTGTAGGTATGCG | R |
| *HvDLO1-like* | Horvu_MOREX_2H01G572800 | LOC123430617 | TCATGGTGACGAACCACGG | F |
| | | | CTTGAGCCGTTCAGACTCCG | R |
| *HvbHLH167-like* | Horvu_MOREX_3H01G001000 | LOC123442338 | AGAAGACCTCTGTGCAAGCC | F |
| | | | GTGGGTGATGCGTCCAAATAC | R |
| *HvABCG1-like* | Horvu_MOREX_5H01G408900 | LOC123395611 | TACTACTGGCTCCGCTTTGC | F |
| | | | AGGCACCGACGAACATAAGC | R |
| *HvJOX2/4-like* | Horvu_MOREX_4H01G310300 | LOC123448920 | GCCTAAACATCCCGGTAGTGG | F |
| | | | GTTCACCGCCTGGAAGAAAC | R |
| *HvGDSL-like* | Horvu_MOREX_7H01G720100 | LOC123412129 | GCTACCCAAGCAAGGAGGAG | F |
| | | | CTTGACGACGTATGCCTGTG | R |
| *HvPR5* | | LOC123409670 | ACAACTGCGGCTCCACAATA | F |
| | | | GAGAGAGCATGAGAGCTGGC | R |
| *HvEF1α* | | | GTCATTGATGCTCCTGGTCA | F |
| | | | CTGCTTCACACCAAGAGTGA | R |
| *HvUbiquitin* | | | ACCCTCGCCGACTACAACAT | F |
| | | | CAGTAGTGGCGGTCGAAGTG | R |

**Supplementary Table S1** Oligonucleotides used for qPCR. Gene names are derived from BLAST analysis of protein sequences associated with the indicated loci. Locus ID allows comparison to Supplementary Table S2 and Locus (Ensembl Plants) indicates the corresponding locus in the current *Hordeum vulgare* reference genome sequence. Abbreviations: F, forward; R, reverse

| ASV | Genus | $Log_2$-fold Change | *p*-value |
|---|---|---|---|
| **ASV74** | Azohydromonas | -12.20 | <0.05 |
| **ASV97** | Azohydromonas | -10.75 | <0.0005 |
| **ASV118** | Serratia | 30.00 | <0.0005 |
| **ASV221** | Flavobacterium | -10.30 | <0.05 |
| **ASV272** | Flexivirga | -21.62 | <0.0005 |
| **ASV316** | Kocuria | 29.62 | <0.0005 |
| **ASV328** | Porphyromonas | -21.24 | <0.0005 |
| **ASV340** | Acidovorax | -24.29 | <0.0005 |
| **ASV358** | Intrasporangium | -22.99 | <0.0005 |
| **ASV424** | Porphyromonas | -20.15 | <0.0005 |
| **ASV460** | Corynebacterium | -21.20 | <0.0005 |
| **ASV466** | Order: Solibubrobacterales | 27.54 | <0.0005 |
| **ASV553** | Bacillus | -20.21 | <0.0005 |
| **ASV568** | Alteribacillus | 21.01 | <0.0005 |
| **ASV586** | Streptomyces | -24.69 | <0.0005 |
| **ASV605** | Truepera | -20.12 | <0.0005 |
| **ASV620** | Order: Ktedonobacterales | -17.58 | <0.0005 |

**Supplementary Table S4** Relative abundance of distinct ASVs analysed by 16S rRNA gene amplicon sequencing in samples from WCS417r-treated as compared to control-treated barley plants. Data are derived from six biologically independent replicate experiments. ASVs were assigned to a genus or order by using NCBI BLAST. $Log_2$-fold change between WCS417r- and control-treated plants was calculated using the R package DESeq2, and significance (p-value) was calculated using the built-in Wald-test with FDR correction following the Benjamini-Hochberg procedure.

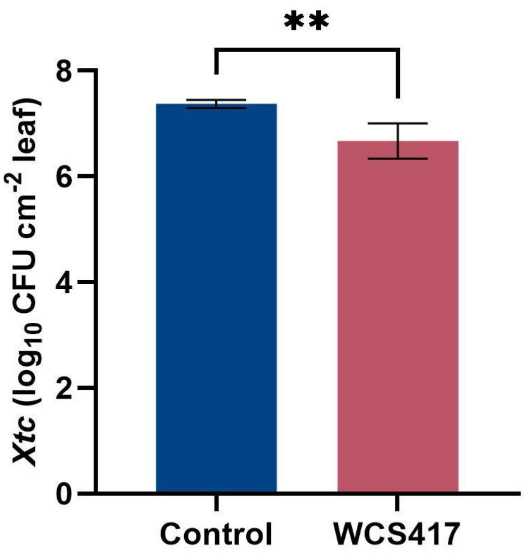


**Supplementary Figure S1** WCS417r-IR against *Xanthomonas translucens* pathovar *cerealis* (*Xtc*). Sterile-grown barley seedlings were treated with *Pseudomonas simiae* WCS417r (WCS417) or an appropriate control solution and subsequently transferred to soil. Three weeks later, leaves were syringe-infiltrated with *Xtc*, colony forming units (CFUs) of which were determined per $cm^2$ of leaf at four days post inoculation (dpi). Bars represent mean $CFUs/cm^2$ +/- SEM from three replicates; asterisks indicate a significant difference (unpaired Student's *t*-test; **, $p < 0.01$). This experiment was repeated seven times; 2 replicates displayed a significant difference between treatments.

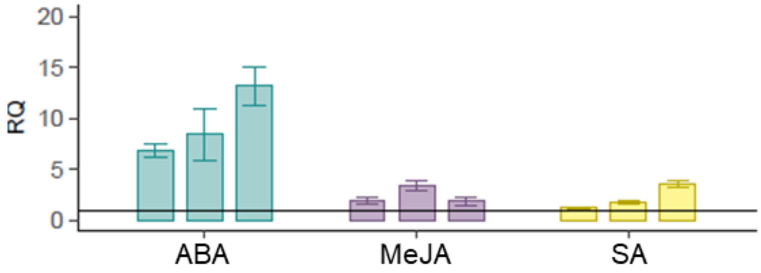


**Supplementary Figure S2** Relative transcript accumulation of *HvPR1* in response to phytohormones. Relative transcript accumulation of *HvPR1* 24 hours post-treatment of barley leaves with abscisic acid (ABA), methyl jasmonate (MeJA), or salicylic acid (SA), as indicated below the panel. Transcript accumulation was normalized to that of *HvEF1α* and is shown relative to the appropriate controls indicated by a black line. Bars represent average values +/- SD from three technical replicates; the results from three biologically independent replicate experiments are shown in individual bars for each treatment.

| Treatment | No of DEGs |
|---|---|
| SA | 149 |
| MeJa | 695 |
| ABA | 116 |

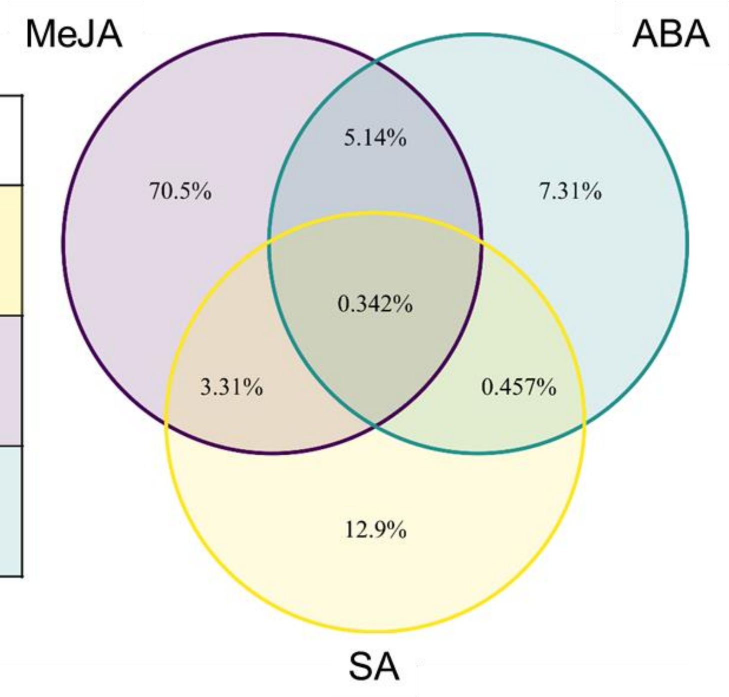


**Supplementary Figure S3** VENN diagram of phytohormone-responsive differentially expressed genes (DEGs) in barley. The data summarize DEGs detected by RNA-sequencing 24 hours post treatment of barley leaves with 100 µM MeJA, ABA, or SA as compared to water-treated controls. Data are derived from three biologically independent replicate experiments. The number of DEGs detected per treatment is indicated in the Table (left) and the percentage of overlapping DEGs between treatments is indicated in the Venn diagram (right).

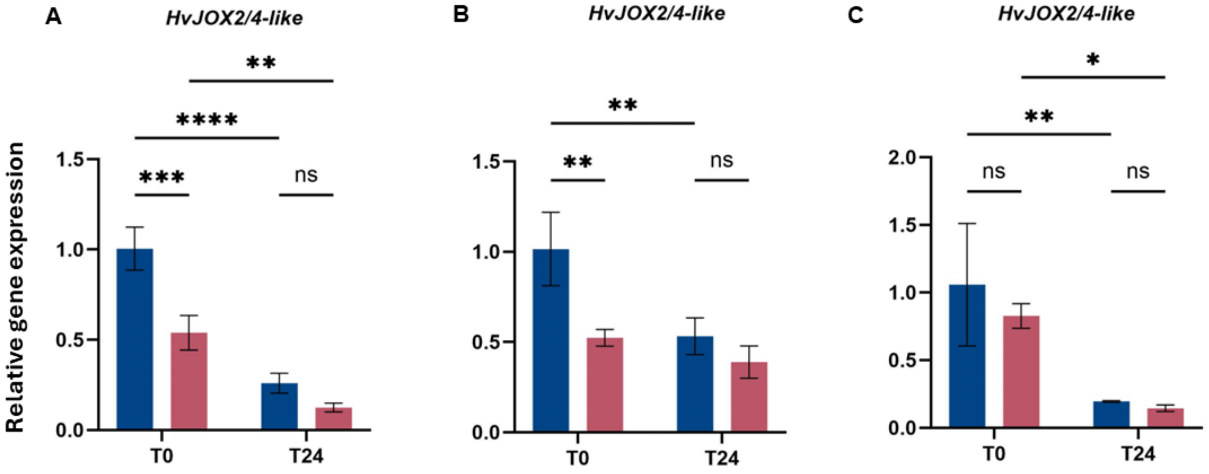


**Supplementary Figure S4** *HvJOX2/4-like* transcript accumulation during WCS417-IR against *Bgh*. *HvJOX2/4-like* transcript accumulation was analysed in WCS417r-treated (indicated in red) as compared to control-treated barley plants (indicated in blue) before (T0) and 24 h (T24) after infection of the plants with *Bgh*. Transcript accumulation was normalized to that of *HvEF1α* and is indicated relative to the control at T0. Panels (**A**), (**B**), and (**C**) represent the results of three biologically independent experiments, including three technical replicates each. Bars represent mean values +/- SD; asterisks indicate level of significance (Tukey's HSD test *, $p < 0.05$, **, $p < 0.01$, ***, $p < 0.001$, ****, $p < 0.0001$; ns, not significantly different).

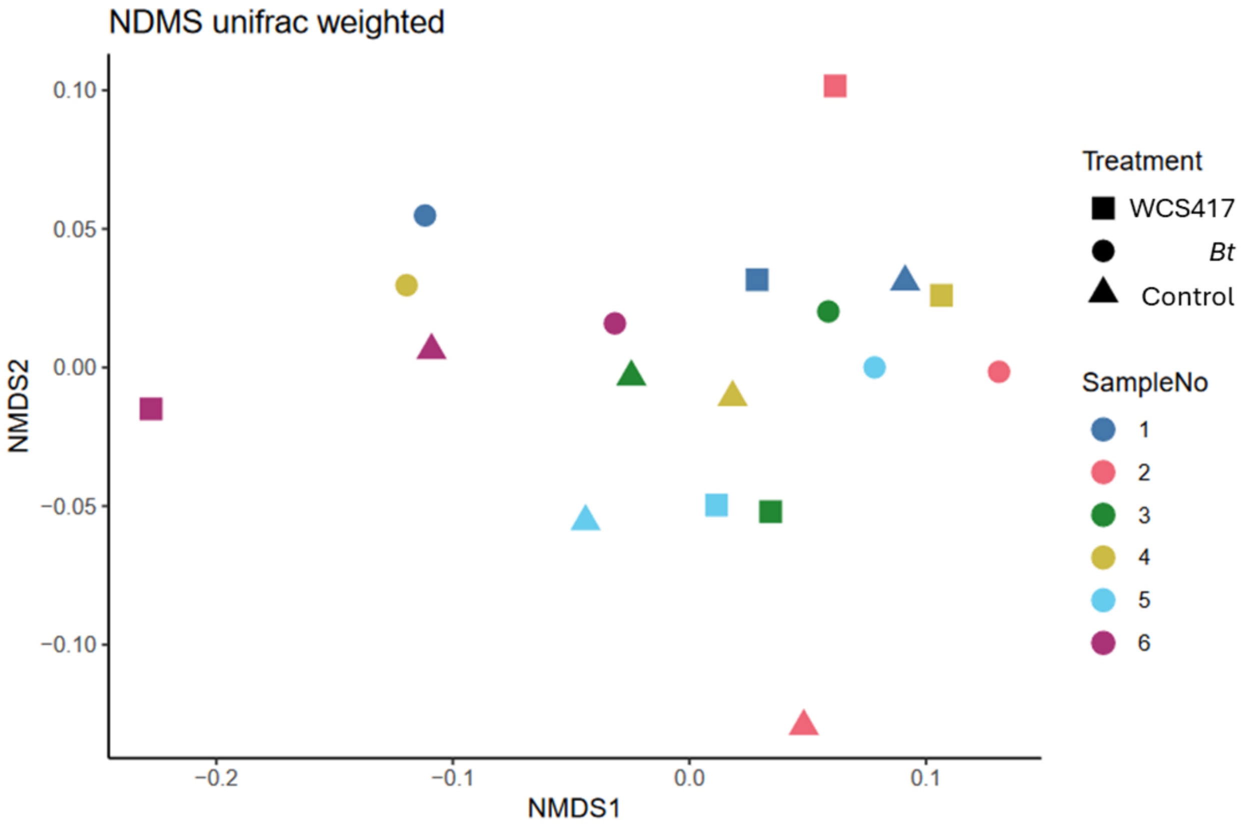


**Supplementary Figure S5** Nonmetric multidimensional scaling **(**NMDS) of microbial composition per treatment and replicate. 16S rRNA gene amplicon sequencing data was analysed by NMDS as calculated from weighted Unifrac distances. Rectangles represent WCS417r-treated plants, circles represent *Bt*-treated plants, and triangles represent the control-treatment. Colours indicate the different independent biological replicates as indicated on the right.

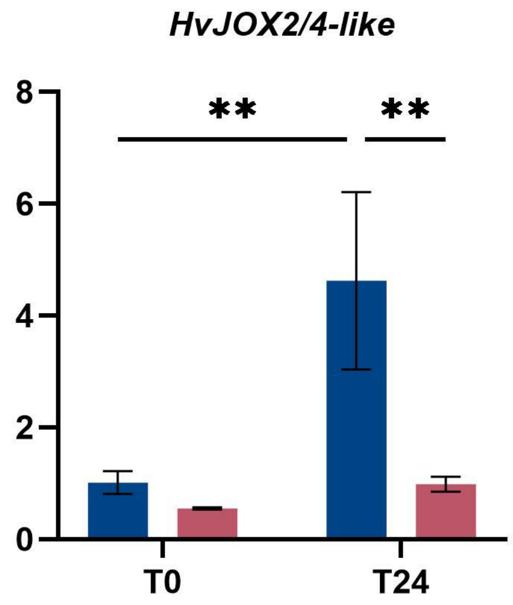


**Supplementary Figure S6** *HvJOX2/4-like* transcript accumulation in response to *Xtc* in plants undergoing WCS417r-IR. *HvJOX2/4-like* transcript accumulation was analysed in WCS417r-treated (indicated in red) as compared to control-treated barley plants (indicated in blue) before (T0) and 24 h (T24) after infection of the plants with *Xtc.* Transcript accumulation was normalized to that of *HvEF1α* and is indicated relative to the control at T0. Bars represent mean values +/- SD from a representative experiment, which was repeated three times with comparable results. Asterisks indicate level of significance (Tukey's HSD test **, p <0.01).
